# Kinesin-1 autoinhibition facilitates the initiation of dynein cargo transport

**DOI:** 10.1101/2022.05.30.493994

**Authors:** Rongde Qiu, Jun Zhang, Xin Xiang

**Affiliations:** Department of Biochemistry and Molecular Biology, The Uniformed Services University of the Health Sciences- F. Edward Hébert School of Medicine, Bethesda, Maryland 20814, USA

**Keywords:** Kinesin-1 autoinhibition, dynein, phi mutation, cargo adapter, *Aspergillus nidulans*, genetic screen

## Abstract

Kinesin-1 undergoes autoinhibition but its functional significance has been unclear. Kinesin-1 transports multiple cargoes including cytoplasmic dynein to the microtubule plus ends. From a genetic screen for *Aspergills* mutants defective in dynein-mediated early endosome transport, we identified a kinesin-1 mutation *kinA*^K895*^ that disrupts kinesin-1 autoinhibition. Consistent with *kinA*^K895*^ making kinesin-1 constitutively active, the mutant proteins accumulate abnormally near the microtubule plus ends. Unexpectedly, our genetic data show that kinesin-1 autoinhibition is unnecessary for transporting its cargoes such as secretory vesicles. Dynein accumulates normally at the microtubule plus ends in the *kinA*^K895*^ mutant. However, the frequency but not the speed of dynein-mediated early endosome transport is significantly decreased, indicating that kinesin-1 autoinhibition facilitates dynein to initiate its cargo transport. Furthermore, kinesin-1 autoinhibition promotes dynein cargo initiation in a way mechanistically distinct from LIS1-promoted dynein switching from its autoinhibited form. Thus, while dynein activation involves dynactin, cargo adapter and LIS1, this study adds kinesin-1 autoinhibition as a new regulatory factor in vivo.

## Introduction

Kinesin-1s are plus-end-directed microtubule motors that transport a variety of cellular cargoes, and defects or dysregulation of kinesin-1s have been implicated in many human diseases (Boecker et al., 2021; Brenner et al., 2018; Gennerich and Vale, 2009; Hirokawa and Tanaka, 2015; Kelliher et al., 2018; Keren-Kaplan and Bonifacino, 2021; Kruppa and Buss, 2021; Nicolas et al., 2018; Roney et al., 2021). The activity of kinesin-1 is regulated by multiple factors, including its light chains, cargo adapter proteins, an opposite motor that binds to the same cargo, phosphorylation, and microtubule-binding proteins (Ally et al., 2009; Blasius et al., 2007; Byrd et al., 2001; Chen and Sheng, 2013; Coy et al., 1999; Fenton et al., 2021; Ferro et al., 2022; Fu and Holzbaur, 2013; Gindhart et al., 1998; Guardia et al., 2021; Keren-Kaplan and Bonifacino, 2021; Monroy et al., 2020; Sakamoto et al., 2005; Twelvetrees et al., 2019; Verhey and Hammond, 2009; Williams et al., 2014; Xu et al., 2012; Zhao et al., 2021). An important aspect of kinesin-1 regulation is its autoinhibition that involves the interaction between the C-terminal tail and the motor domains (Cai et al., 2007; Chiba et al., 2022; Coy et al., 1999; Dietrich et al., 2008; Friedman and Vale, 1999; Kaan et al., 2011; Seiler et al., 2000; Stock et al., 1999; Verhey and Hammond, 2009; Wong et al., 2009). Recently, an exon27-skipping mutation (Δexon27) in the KIF5A kinesin-1 that causes the devastating neurodegenerative disease Amyotrophic Lateral Sclerosis (ALS) has been linked to a loss of autoinhibition (Baron et al., 2022; Nakano et al., 2022; Pant et al., 2022). However, the functional significance of kinesin-1 autoinhibition in normal cells would need to be further dissected.

Cytoplasmic dynein is the minus-end-directed microtubule motor, and defects in dynein and/or its regulator dynactin have been implicated in ALS (Chevalier-Larsen and Holzbaur, 2006; LaMonte et al., 2002; Maimon et al., 2021; Münch et al., 2004). Some vesicular cargos bind both dynein and kinesin-1 via the same cargo adapter (Cox and Spradling, 2006; Fenton et al., 2021; Fu and Holzbaur, 2014)(Canty et al., 2021, bioRxiv; Celestino et al., 2021, bioRxiv), and in this case, loss of kinesin-1 autoinhibition may help overcome the force of dynein to change cargo distribution (Baron et al., 2022; Kelliher et al., 2018). Interestingly, in many cell types including filamentous fungi, mammalian neurons, *C. elegans* neurons, and Drosophila oocytes, dynein itself is a cargo of kinesin-1 (Arimoto et al., 2011; Brendza et al., 2002; Duncan and Warrior, 2002; Egan et al., 2012; Hirokawa et al., 1990; Januschke et al., 2002; Lenz et al., 2006; Twelvetrees et al., 2016; Yamada et al., 2010; Zhang et al., 2003). In mammalian neurons, kinesin-1 interacts with dynein directly via kinesin light chains and moves dynein towards the microtubule plus ends (Ligon et al., 2004; Twelvetrees et al., 2016). In filamentous fungi, kinesin-1s are largely (although not completely) responsible for the microtubule plus-end accumulation of dynein and its regulator dynactin (Egan et al., 2012; Lenz et al., 2006; Penalva et al., 2017; Yao et al., 2012; Zhang et al., 2003; Zhang et al., 2010). Fungal kinesin-1s do not have associated light chains (Seiler et al., 2000; Steinberg and Schliwa, 1995), but dynactin is important for the dynein-kinesin-1 interaction (Qiu et al., 2018). The kinesin-1-mediated dynein accumulation at the plus ends is important for the retrograde transport of dynein cargoes such as early endosomes, most likely because it enhances the chance for dynein-cargo interaction (Abenza et al., 2009; Egan et al., 2012; Lenz et al., 2006; Zekert and Fischer, 2009; Zhang et al., 2010). It is important to note that the plus-end-directed transport of early endosomes does not use kinesin-1 but uses kinesin-3 (Lenz et al., 2006; Wedlich-Soldner et al., 2002; Zekert and Fischer, 2009). Dynein transports early endosomes away from the plus ends with the help of dynactin and the FTS-Hook-FHIP complex (Bielska et al., 2014; Yao et al., 2014; Zhang et al., 2014; Zhang et al., 2011b). The mechanism of dynein-mediated early endosome transport is conserved and has been further dissected using mammalian proteins (Christensen et al., 2021; Guo et al., 2016; Lau et al., 2021; Olenick et al., 2019; Schroeder and Vale, 2016; Urnavicius et al., 2018; Yeh et al., 2012), but factors regulating this process need to be further studied. In the case of kinesin-1, it was unknown whether its autoinhibition plays a role in the initiation of dynein cargo transport from the microtubule plus ends. During a genetic screen for mutants defective in dynein-mediated early-endosome transport in *Aspergillus nidulans*, we serendipitously identified a nonsense mutation *kinA*^K895*^ causing a loss of kinesin-1 autoinhibition. Our analyses suggest that this autoinhibition is unnecessary for kinesin-1 to transport its own cargos including dynein and secretory vesicles, but it is important for the initiation of dynein-mediated early endosome transport.

## Results and Discussion

### Identifying *kinA*^K895*^ as a causal mutation affecting dynein-mediated early endosome transport

In *A. nidulans*, the plus ends of microtubules face the hyphal tip and minus ends are either at the spindle-pole body or at septum (Efimov et al., 2006; Egan et al., 2012; Gao et al., 2019; Han et al., 2001; Konzack et al., 2005; Oakley et al., 1990; Xiong and Oakley, 2009; Zeng et al., 2014; Zhang et al., 2017b). Dynein normally accumulates at the microtubule plus ends near the hyphal tip, as represented by the comet-like structures (Han et al., 2001; Xiang et al., 2000). A defect in dynein-mediated early endosome transport causes an abnormal accumulation of early endosomes and their hitchhiking cargoes near the hyphal tip (Abenza et al., 2009; Egan et al., 2012; Lenz et al., 2006; Salogiannis and Reck-Peterson, 2017; Zekert and Fischer, 2009; Zhang et al., 2011b; Zhang et al., 2010). To find additional dynein regulators, we performed genetic screens for early-endosome distribution (*eed*) mutants, which have allowed us to find novel dynein regulators (Xiang and Qiu, 2020). In this study, we performed UV mutagenesis using a strain containing mCherry-labeled RabA and GFP-labeled dynein heavy chain, which allows observation of early endosomes and dynein respectively (Pinar and Peñalva, 2021; Zhuang et al., 2007). The *eedE*16 mutant isolated after UV mutagenesis formed a colony smaller than that of a wild-type strain (Figure 1A). In the mutant, GFP-dynein formed the plus-end comets (Figure 1B), but early endosomes abnormally accumulated near the hyphal tip (Figure 1B).

**Figure 1.**
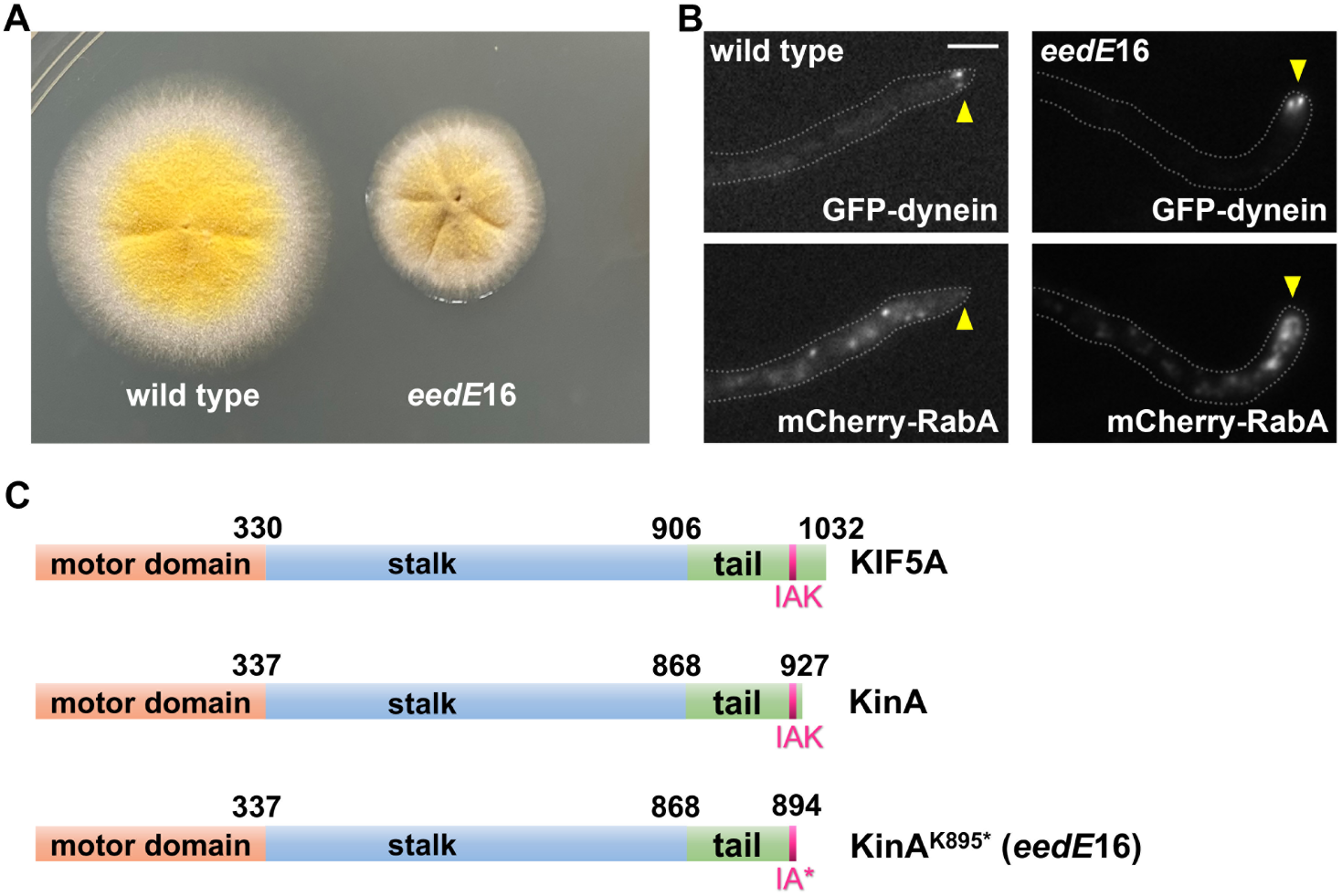
Phenotype of the *eedE*16 mutant and position of the causal mutation in kinesin-1. (A) Colony phenotypes of the *eedE*16 mutant and a wild-type strain. (B) Microscopic images showing the distributions of GFP-dynein and mCherry-RabA-labeled early endosomes (mCherry-RabA) in wild type and the mutant. Bar, 5 μm. Although bi-directional movements of mCherry-RabA-labeled early endosomes are not completely abolished, most of the *eedE*16 hyphal tips (∼80%) show an obvious accumulation of mCherry-RabA signals (n=130). Hyphal tip is indicated by a yellow arrowhead. (C) Domain structures of kinesin-1 proteins in human (KIF5A) and *A. nidulans* (KinA). Positions of the IAK motif involved in kinesin-1 autoinhibition and the *eed*E16 mutation *kinA*^K895*^ within the IAK motif are indicated.

To identify the *eedE*16 mutation, we took a whole-genome sequencing approach as described previously (Qiu et al., 2020). Whole-genome sequencing was done using the genome sequencing and bioinformatic service of Genewiz (www.genewiz.com). We used the software Integrative Genomics Viewer (IGV 2.4.3) to visualize the genomic sequencing data to identify the mutation. The *eedE16* mutation was found in *kinA*, which encodes the only kinesin-1 in *A. nidulans*, with 927 amino acids (Requena et al., 2001). The mutation substituted an A with T that changed the codon for lysine (K) (AAG) to a stop codon (TAG) at residue 895. This results in the deletion of 33 amino acids at the C-terminus of KinA. The C-terminus of Drosophila Kinesin-1 or a human kinesin-1 KIF5A contains two important regions: the IAK motif involved in autoinhibition (Kaan et al., 2011), and the RKRYQ region responsible for ATP-independent microtubule binding (Lu et al., 2016; Winding et al., 2016). The IAK motif involved in autoinhibition is conserved in KinA but the residues for ATP-independent microtubule binding are not (Supplemental Figure 1). Importantly, K895 corresponds to K within the IAK motif (Figure 1C; Supplemental Figure 1), and the mutation should delete it and the rest of the C-terminus. Thus, the *kinA*^K895*^ mutation is predicted to eliminate kinesin-1 autoinhibition.

### KinA^(1-894)^-GFP forms an abnormally strong accumulation near the microtubule plus ends

To compare the behaviors of the wild-type and the *kinA*^K895*^ mutant kinesin-1s, we constructed strains with either the *kinA*-GFP or the *kinA*^(1-894)^-GFP fusion gene integrated at the endogenous *kinA* locus (Figure 2A). KinA-GFP formed a diffuse background in the cytoplasm with only very faint plus-end comet-like structures near the hyphal tip (Figure 2B). In contrast, KinA^(1-894)^-GFP signals were strongly accumulated at the hyphal tip near microtubule plus ends. This is consistent with the localization pattern of uninhibited kinesin-1s in neurons and also in another fungus *Neurospora crassa* (Baron et al., 2022; Seiler et al., 2000; Twelvetrees et al., 2016).

**Figure 2.**
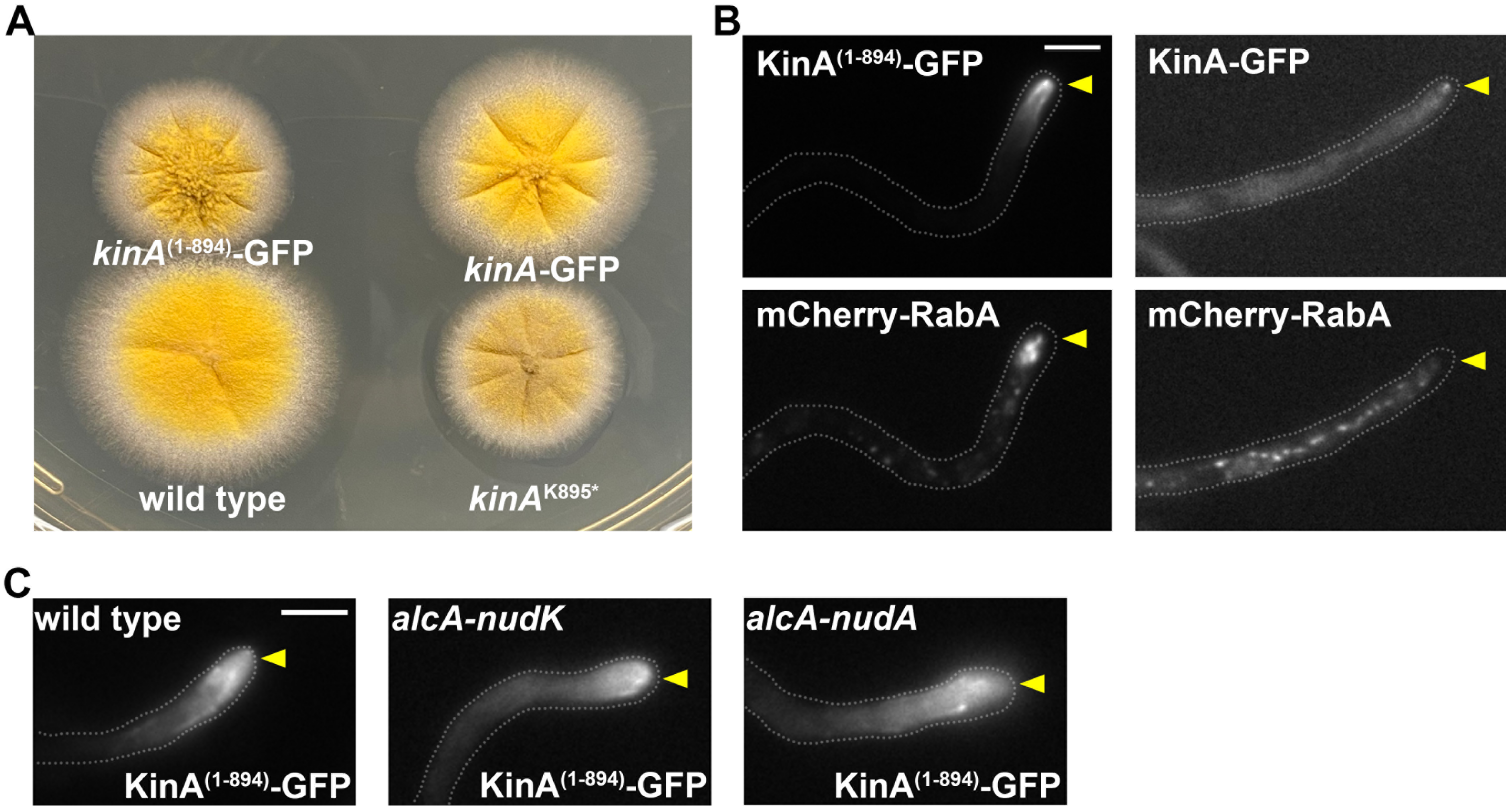
KinA^(1-894)^-GFP fusion proteins form a prominent accumulation near the microtubule plus ends. (A) Colony phenotypes of the strains containing *kinA*-GFP and *kinA*^(1-894)^-GFP in comparison to wild type and the *eed*E16 mutant *kinA*^K895^*. (B) Localization of KinA-GFP or KinA^(1-894)^-GFP and mCherry-RabA-labeled early endosomes in strains containing one of the GFP fusions. Most (∼87.5%) of the hyphal tips of the *kinA*^(1-894)^-GFP mutant show an abnormal accumulation of early endosomes (n=32). Hyphal tip is indicated by a yellow arrowhead. Bar, 5 μm. (C) The hyphal-tip localization of KinA^(1-894)^-GFP is independent of dynein or dynactin. Images were taken from cells grown on glucose, a repressive medium for the regulatable *alcA* promoter. The *alcA-nudK*^Arp1^ or the *alcA-nudA*^HC^ conditional-null allele allows the expression of dynactin Arp1 or dynein heavy chain to be shut off on glucose. Hyphal tip is indicated by a yellow arrowhead. Bar, 5 μm.

As expected, early endosomes (labeled with mCherry-RabA) distributed normally in the kinA-GFP strain but accumulated abnormally near the hyphal tip in the kinA^(1-894)^-GFP strain just like in the *kinA*^K895*^ mutant (Figure 2B). Using strains containing the conditional-null allele of dynactin Arp1 (*alcA-nudK*^Arp1^) or dynein heavy chain (*alcA-nudA*^HC^), we further showed that the prominent hyphal-tip accumulation of KinA^(1-894)^-GFP signals occurs in the absence of dynein or dynactin (Figure 2C).

### Kinesin-1 autoinhibition is not critical for the transport of its own c argo

We next sought to determine if kinesin-1 auto-inhibition is critical for its cargo-transporting function. To do that, we compared the effect caused by the C-terminal truncated *kinA* with that of the previously constructed Δ*kinA* allele (Requena et al., 2001). The colony formed by the *kinA*^K895*^ mutant is bigger than the Δ*kinA* mutant (Figure 3A, 3B), suggesting that function of kinesin-1 is partially retained in the uninhibited mutant. In filamentous fungi, both kinesin-1 and myosin-V are capable of transporting secretory vesicles to support hyphal tip extension (Pantazopoulou et al., 2014; Penalva et al., 2017; Schuchardt et al., 2005; Schuster et al., 2012). In *A. nidulans*, the Δ*kinA* or the *alcA*-*myoV* (myosin-V conditional-null mutant) single mutant forms a colony whose size is smaller than that of wild type, but the Δ*kinA, alcA*-myoV double mutant is nearly inviable when grown on glucose that shuts off the expression of myosin-V (Zhang et al., 2011a) (Figure 3C). In contrast, the colony of the *kinA*^(1-894)^-GFP, *alcA*-*myoV* double mutant was slightly bigger than the *alcA*-*myoV* single mutant (Figure 3C, 3D). To confirm this result, we also made a *kinA*^(1-894)^-GFP, Δ*myoV* double mutant. The Δ*myoV* single mutant formed a compact colony (Taheri-Talesh et al., 2012). In striking contrast to the nearly lethal phenotype of the Δ*kinA*, Δ*myoV* double mutant (Penalva et al., 2017), the *kinA*^(1-894)^-GFP, Δ*myoV* double mutant formed a colony whose diameter is similar to that of the Δ*myoV* single mutant (Figure 3C, 3D). These genetic data strongly indicate that auto-inhibition is not necessary for the general transporting function of kinesin-1 involved in supporting hyphal growth. Consistent with this conclusion, Rab11-marked secretory vesicles, which are cargoes of kinesin-1 and myosin-V (Pantazopoulou et al., 2014; Penalva et al., 2017; Pinar et al., 2022), reach the hyphal tip in the *kinA*^K895*^, Δ*myoV* double mutant just like in the Δ*myoV* single mutant (Figure 3E).

**Figure 3.**
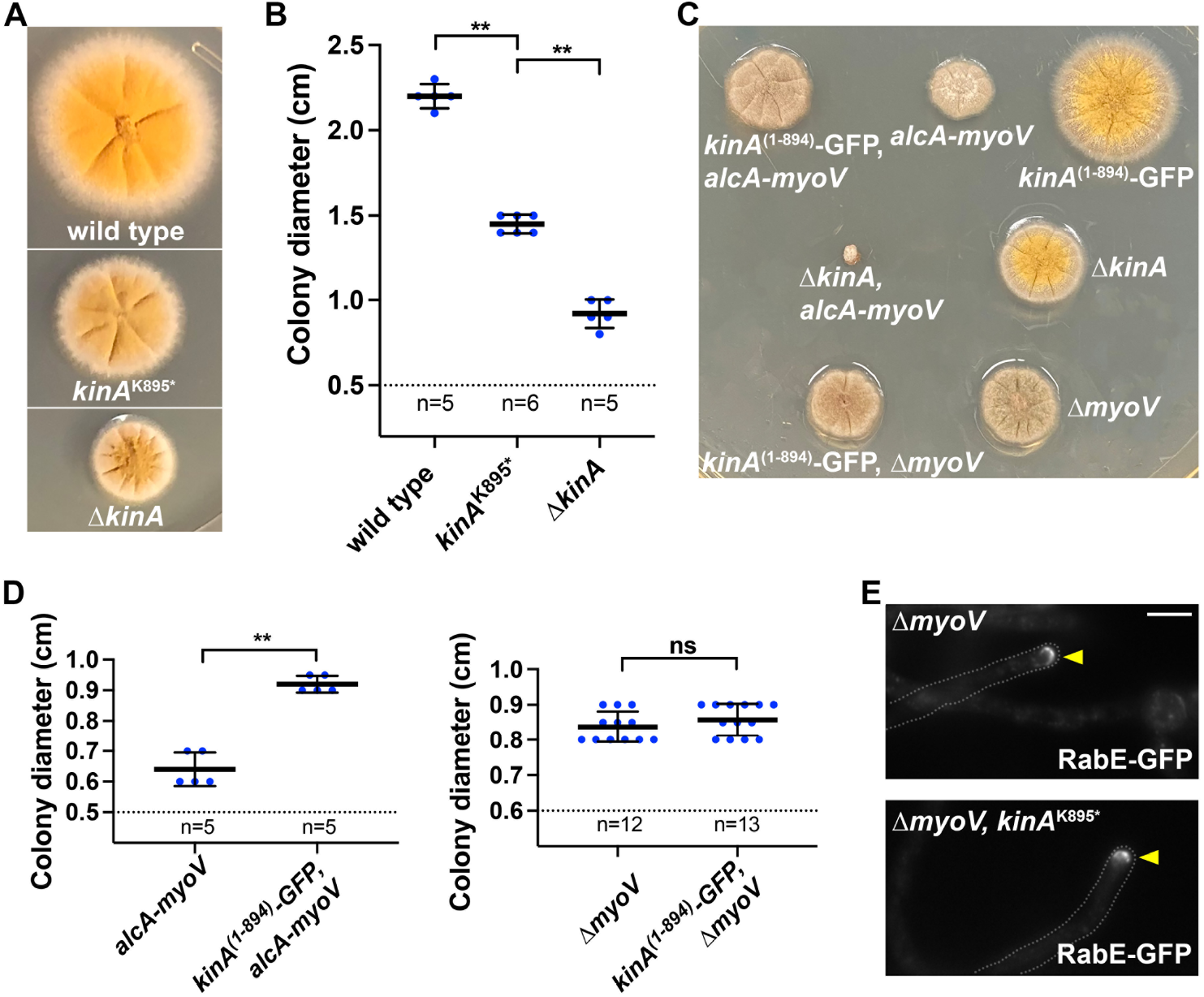
Kinesin-1 autoinhibition is not critical for its own cargo transport function. (A) Colony phenotypes of a wild-type strain, the *kinA*^K895*^ mutant and the Δ*kinA* mutant. (B) A quantitative analysis of colony diameter of the wild-type strain, the *kinA*^K895*^ mutant and the Δ*kinA* mutant. Scatter plots with mean and S.D. values were generated by Prism 9. The Mann-Whitney test (unpaired, two tailed) was used for analyzing two data sets at a time without assuming normal distribution of the data. **p<0.01. (C) Colony phenotypes of the single and double myosin-V (*myoV*) and kinesin-1 (*kinA*) mutants. (D) A quantitative analyses of colony diameters of *alcA-myoV* and *kinA*^(1-894)^-GFP, *alcA-myoV* (**p<0.01; unpaired, Mann-Whitney test) and of Δ*myoV* and *kinA*^(1-894)^-GFP, Δ*myoV*. The difference between the Δ*myoV* and *kinA*^(1-894)^-GFP, Δ*myoV* strains is not significant (ns) at p=0.05 (unpaired, Mann-Whitney test). (E) Microscopic images showing the hyphal-tip signals of RabE-GFP in Δ*myoV* and the Δ*myoV, kinA*^K895^* double mutant. Hyphal tip is indicated by a yellow arrowhead. Bar, 5 μm.

### Loss of kinesin-1 autoinhibition affects the initiation of dynein-mediated early endosome transport

We next quantitated the microtubule plus-end dynein localization in the *kinA*^K895*^ mutant, and we did this in the Δ*hookA* background missing the cargo adapter (Note that cargo adapters activate dynein to leave the plus ends (Baumbach et al., 2017; Jha et al., 2017; Lammers and Markus, 2015; McKenney et al., 2014; Qiu et al., 2019; Schlager et al., 2014; Splinter et al., 2012)). In the *kinA*^K895*^, Δ*hookA* double mutant, the plus-end dynein comet intensity is similar to that in the Δ*hookA* single mutant (Figure 4A, 4B). This is consistent with a previous study showing that the posterior localization of dynein during Drosophila oogenesis needs kinesin-1 but not its IAK region implicated in autoinhibition (Williams et al., 2014). Thus, although uninhibited KinA tends to move to the microtubule plus ends without carrying its cargo, it can still carry dynein to the plus end. This may be explained by in vitro data showing that an uninhibited kinesin-1 is more likely to land on a microtubule (Baron et al., 2022; Chiba et al., 2022) or have a higher processivity (Friedman and Vale, 1999; Pant et al., 2022) than a normally autoinhibited kinesin-1.

**Figure 4.**
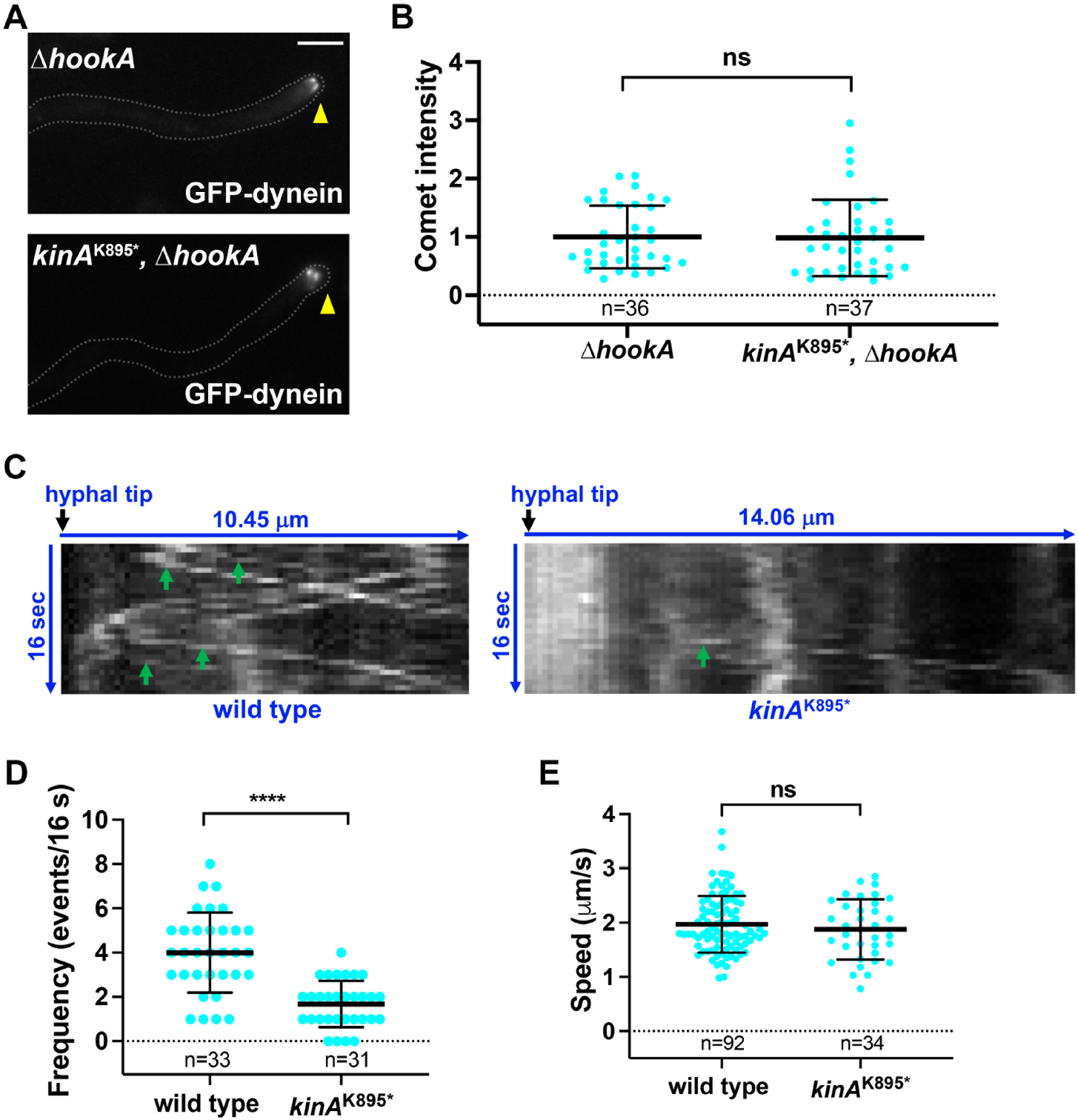
Loss of kinesin-1 autoinhibition does not affect dynein accumulation at the microtubule plus ends but affects the initiation of dynein-mediated early endosome transport. (A) Microscopic images showing the hyphal tip signals of GFP-dynein in the Δ*hookA* single and the *kinA*^K895^*, Δ*hookA* double mutant. Hyphal tip is indicated by a yellow arrowhead. Bar, 5 μm. (B) A quantitative analysis on GFP-dynein comet intensity in Δ*hookA* and the Δ*hookA, kinA*^K895^* double mutant. All values are relative to the average value for Δ*hookA*, which is set as 1. Scatter plots with mean and S.D. values were generated by Prism 9. The Mann-Whitney test (unpaired, two tailed) was used for analyzing the two data sets without assuming normal distribution of the data. (C) Kymographs showing the movements of early endosomes in wild type and the *kinA*^K895*^ mutant. Green arrows indicate dynein-mediated movements away from the hyphal tip. (D) A quantitative analysis on the frequency of dynein-mediated transport in wild type (n=33 hyphal tips) and the *kinA*^K895*^ mutant (n=31 hyphal tips). Scatter plots with mean and SD values were generated by Prism 9. ****p<0.0001 (unpaired, Mann-Whitney test, Prism 9). (E) A quantitative analysis on the speed of dynein-mediated early endosome movement in wild type (n=92 movements) and the *kinA*^K895*^ mutant (n=34 movements). The difference between wild type and the mutant is not significant (ns) at p=0.05 (unpaired, Mann-Whitney test, Prism 9).

To better understand the effect of kinesin-1 autoinhibition on dynein function, we measured the frequency and speed of dynein-mediated early endosome movement in *kinA*^K895*^ mutant in comparison to the wild-type control. We found that the frequency but not the speed is significantly reduced in the mutant (Figure 4C, 4D, 4E). Thus, although dynein can be targeted to the microtubule plus ends in the *kinA*^K895*^ mutant, its function in initiating cargo transport is compromised. This defect is conceptually similar to (although less severe than) that caused by loss of LIS1 (Markus et al., 2020), a dynein regulator required for the initiation of dynein-mediated early endosome transport in filamentous fungi (Egan et al., 2012; Lenz et al., 2006).

### The role of kinesin-1 autoinhibition in dynein cargo initiation differs from that of LIS1

Dynein activation involves conformational changes of the dynein heavy chain dimer from an auto-inhibited “phi” conformation to the open conformation (Torisawa et al., 2014; Zhang et al., 2017a), and binding of dynactin and cargo adapter to the dynein tail causes dynein to assume a parallel conformation allowing it to move processively (Zhang et al., 2017a). The phi mutation keeps dynein open, thereby facilitating its activation and centrosomal accumulation (Zhang et al., 2017a). In *A. nidulans*, the phi mutation (*nudA*^R1602,K1645E^) or overexpression of the ΔC-HookA adapter causes dynein to relocate from the microtubule plus ends to the minus ends on septa or spindle-pole bodies (Qiu et al., 2019). Since the spindle-pole body accumulation of activated dynein is cell-cycle dependent (Bieger et al., 2021), we only focus on the septal minus ends here (Zhang et al., 2017b). Upon loss of LIS1 function in *nudF* mutants (Xiang et al., 1995), ΔC-HookA overexpression fails to drive the relocation to septa but the phi mutation is able to do so, suggesting that a main function of NudF/LIS1 is to overcome the autoinhibited phi conformation (Qiu et al., 2019). This notion is consistent with data from other groups especially in budding yeast (Elshenawy et al., 2020; Gillies et al., 2022; Htet et al., 2020; Marzo et al., 2020). In the *kinA*^K895*^ mutant, either the phi mutation or overexpression of the ΔC-HookA adapter can still cause dynein relocation to septa like in cells containing the wild-type *kinA* allele, but in either case the relocation is partially defective as evidenced by the more easily observable plus-end dynein comets and the reduced septal signals (Figure 5A, 5B, 5C, 5D). Importantly, quantitation of septal dynein signals indicates that overexpression of ΔC-HookA is more effective than the phi mutation in driving dynein relocation in the *kinA*^K895*^ mutant (Figure 5B, 5C, 5D), which is opposite to the situation in the *nudF* (lis1) mutants. Furthermore, the phi mutation significantly reduces the hyphal-tip early endosome accumulation in the *nudF*6 mutant but not in the *kinA*^K895*^ mutant (Figure 5E, 5F). Thus, kinesin-1 autoinhibition is involved in the initiation of dynein cargo transport with a mechanism different from that of LIS1-mediated dynein opening.

**Figure 5.**
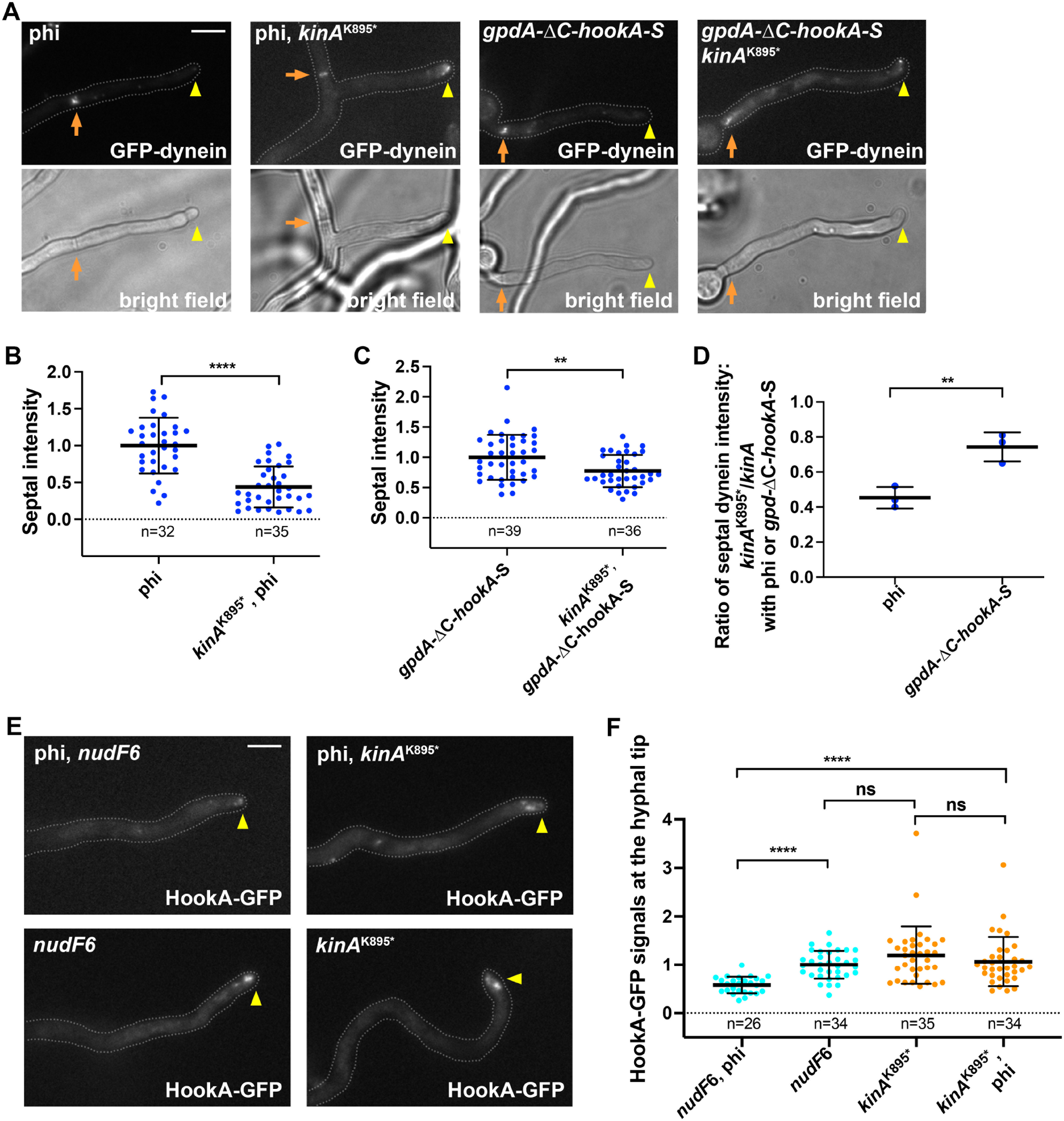
Effects of factors involved in dynein activation on the *kinA*^K895*^ mutant. (A) Dynein localization in the phi single mutant and the phi, *kinA*^K895*^ double mutant, and that upon overexpression of the cargo adapter ΔC-HookA in the wild-type background (*gpdA*-ΔC-*hookA*-S) and in the *kinA*^K895*^ background (*kinA*^K895*^, *gpdA*-ΔC-*hookA*-S). Bright-field images are shown below to indicate hyphal shape and position of septum. Hyphal tip is indicated by a yellow arrowhead and septum by a brown arrow. Bar, 5 μm. (B) Quantitation of dynein signals at septa (for the two phi-mutation-containing strains). All values are relative to the average value for the phi single mutant, which is set as 1. Scatter plots with mean and SD values were generated by Prism 9. ****p<0.0001 (unpaired, Mann-Whitney test, Prism 8). (C) Quantitation of dynein signals at septa (for the two *gpdA*-ΔC-*hookA*-S-containing strains). All values are relative to the average value for the *gpdA*-ΔC-*hookA*-S strain, which is set as 1. Scatter plots with mean and SD values were generated by Prism 9. **p<0.01 (unpaired, Mann-Whitney test, Prism 9). (D). Ratio of the *kinA*^K895*^ mutant value to the wild type value in the phi or the *gpdA*-ΔC-*hookA*-S strains. Data were from three experiments. Scatter plots with mean and SD values were generated by Prism 9. **p<0.01 (unpaired, Mann-Whitney test, Prism 9). (E) Effect of the phi mutation on the hyphal-tip accumulation of early endosomes (labeled by HookA-GFP) in the *nudF*6 (*lis1*) and *kinA*^k895*^ mutants. Images of HookA-GFP in the *nudF*6 single and the *nudF*6, phi double mutant, and in the *kinA*^k895*^ single and the *kinA*^k895*^, phi double mutant. Because we routinely culture the *kinA*^k895*^ mutant at 32°C, we used 32°C for culturing the *nudF*6 mutant in this experiment. Note that although the function of NudF/LIS1 is not completely lost at 32°C (a semi-restrictive temperature), the hyphal-tip accumulation of HookA-GFP signals is very obvious in the single *nudF*6 mutant. Bar, 5 μm. (F) A quantitative analysis on hyphal-tip accumulated HookA-GFP signals. The average value for the *nudF*6 strain is set as 1. Scatter plots with mean and S.D. values were generated by Prism 9. ****p<0.0001; ns, non-significant or p>0.05 (Kruskal-Wallis with Dunn’s multiple comparisons test, unpaired).

How kinesin-1 autoinhibition facilitates the initiation of dynein cargo transport remains an important question. Kinesin-mediated transport of dynein to the microtubule plus ends has been reported from fungi to mammalian neurons (although kinesin-7 instead of kinesin-1 is used in budding yeast) (Arimoto et al., 2011; Brendza et al., 2002; Carvalho et al., 2004; Duncan and Warrior, 2002; Egan et al., 2012; Hirokawa et al., 1990; Januschke et al., 2002; Lenz et al., 2006; Roberts et al., 2014; Twelvetrees et al., 2016; Yamada et al., 2010; Zhang et al., 2003). In several systems, the plus-end accumulation of dynein and its regulator dynactin play an important role in cargo binding or the initiation of cargo transport (Lenz et al., 2006; Lloyd et al., 2012; Markus and Lee, 2011; Moughamian and Holzbaur, 2012; Splinter et al., 2012; Vaughan et al., 2002; Xiang and Qiu, 2020). However, it is not known how kinesin-1 regulation plays a role in preparing dynein-dynactin for cargo binding and transport from the microtubule plus end. From our current data, we speculate that kinesin-1 autoinhibition can happen after LIS1-mediated dynein opening but needs to occur either before or after dynein-cargo interaction to facilitates the interaction and/or cargo-adapter-mediated dynein activation. If kinesin-1 fails to undergo autoinhibition, even if dynein is switched to the open conformation, it will still fail to initiate cargo transport. Thus, kinesin-1 autoinhibition is a factor that cooperates with LIS1 (Markus et al., 2020) (its binding protein NudE) (Garrott et al., 2022)), dynactin and cargo adapters (Olenick and Holzbaur, 2019; Reck-Peterson et al., 2018) to initiate dynein cargo transport in cells where kinesin-1-mediated dynein accumulation at the microtubule plus ends is important for dynein cargo transport.

Recently, multiple studies indicate that an ALS-causing mutation (Δexon27) that changes the C-terminus of KIF5A is a gain-of-function mutation that leads to a loss of autoinhibition (Baron et al., 2022; Nakano et al., 2022; Pant et al., 2022). Since the mutation creates a new C-terminus, it needs to be addressed whether it is the loss of autoinhibition *per se* or the creation of a new C-terminus that leads to ALS. Our analysis on *A. nidulans* diploids has shown that the *kinA*^K895*^ or the *kinA*^(1-894)^-GFP allele is also a gain-of-function mutation that partially affects dynein-mediated early endosome transport in the presence of a wild-type copy of *kinA* (Supplemental Figure 2). As dynein defects have been linked to ALS (Chevalier-Larsen and Holzbaur, 2006; Gershoni-Emek et al., 2016; Hafezparast et al., 2003; Ikenaka et al., 2013; Kieran et al., 2005; LaMonte et al., 2002; Maimon et al., 2021; Mentis et al., 2022; Münch et al., 2004; Stavoe and Holzbaur, 2019; Ström et al., 2008), it is likely that a partial defect in dynein function contributes to ALS caused by the gain-of-function mutations in KIF5A. Future studies will be needed to test this idea.

## Materials and Methods

### *A. nidulans* strains and media

*A. nidulans* strains used in this study are listed in Supplemental Table 1. UV mutagenesis on spores of *A. nidulans* strains was done as previously described (Willins et al., 1995; Xiang et al., 1999). Solid rich medium was made of either YAG (0.5% yeast extract and 2% glucose with 2% agar) or YAG+UU (YAG plus 0.12% uridine and 0.12% uracil). Genetic crosses and diploid construction were done by standard methods. Solid minimal medium containing 1% glucose was used for selecting progeny from a cross and for selecting diploids. For live cells imaging experiments, cells were cultured in liquid minimal medium containing 1% glycerol for overnight at 32°C. All the biochemical analyses (for confirming strain genotypes) and genomic DNA preparation were done using cells grown at 32°C for overnight in liquid YG rich medium (0.5% yeast extract and 2% glucose). For a few experiments using strains containing the *alcA-nudK*^Arp1^ or *alcA-nudA*^HC^ allele, we harvested spores from the solid minimal medium containing 1% glycerol and cultured them in liquid minimal medium containing 1% glucose for imaging analysis.

### Construction of the strain containing the *kinA*-GFP allele at the *kinA* locus

Strains were constructed by using standard procedures used in *A. nidulans* (Nayak et al., 2006; Szewczyk et al., 2006; Yang et al., 2004). For constructing the *kinA*-GFP fusion, we used the following six oligos to amplify genomic DNA from RQ177 (Qiu et al., 2018) that contains the GFP-*AfpyrG* fusion (Yang et al., 2004): K5U (AGTCTTTCAGAGACGCAGGG), BW2 (TCGTC TATCAAAAAACCAACTTGTG), KF3 (CACAAGTTGGTTTTTTGATAGACGAGGAGCTGGTGCAGGCG), BW4 (CCATCTAG ATATCTGCAGGAAGGGGCTGTCTGAGAGGAGGCACTG), BW5 (CCCCTTCCTGCAGATATCTAGATGG) and BW6 (GCTGAAGTTGGTTGATTTGCGG). Specifically, K5U and BW2 were used to amplify the ∼1-kb fragment in the coding region, and BW5 and BW6 were used to amplify the ∼1-kb fragment in the 3’ untranslated region, and KF3 and BW4 were used to amplify the ∼2.7-kb GFP-*AfpyrG* fragment using genomic DNA from the RQ177 strain (Qiu et al., 2018). We then used two oligos, K5U and BW6, for a fusion PCR of the three fragments to generate the ∼4.6-kb *kinA*-GFP-AfpyrG fragment that we used to transform into XY42 containing Δ*nkuA* (Nayak et al., 2006) and mCherry-RabA (Abenza et al., 2009; Qiu et al., 2018; Zhang et al., 2010). The transformants were screened by microscopically observing the GFP signals, and the presence of the *kinA*-GFP fusion was confirmed by western blotting analysis with a polyclonal anti-GFP antibody from Clontech. In addition, we also performed a diagnostic PCR to verify the correct integration using oligos K5UTR (GAACGACCTCACAGACTCA) and AFpyrG3 (GTTGCCAGG TGAGGGTATTT).

### Construction of the strain containing the *kinA*^(1-894)^-GFP allele at the *kinA* locus

For constructing the *kinA*^(1-894)^-GFP strain, we made the *kinA*^(1-^ 894)-GFP-*AfpyrG* fragment by inserting the GFP-*AfpyrG* fragment into the *kinA*^K895*^ mutation site right before the stop codon. The following four oligos were used to make the *kinA*^(1-^ 894)-GFP-*AfpyrG* construct: K5U, K895.R (GCACCAGCTCCA GCGATTCGGGAGCCAG), K895.F (AATCGCTGGAGCTGGTGCAGGCG) and BW6. Specifically, K5U and K895.R were used to amplify a ∼0.9-kb fragment of the 5’ coding region, and K895.F and BW6 were used to amplify the ∼2.7-kb GFP-*AfpyrG* fragment using genomic DNA from the RQ197 strain (containing *kinA*-GFP) as template. We then used two oligos, K5U and BW6, for a fusion PCR to fuse the two fragments to generate the ∼4.5-kb *kinA*^(1-894)^-GFP-*AfpyrG* fragment that we used to transform the XY42 strain and the RQ54 strain. The transformants were screened by microscopically observing the GFP signals and further confirmed by a western blotting analysis with a polyclonal anti-GFP antibody from Clontech. In addition, we also performed a diagnostic PCR to verify the correct integration using oligos K5UTR (GAACGACCTCACAGACTCA) and GFP-5R (CAGTGAAAAGTTCTTCTCCTTTACT).

### Live-cell imaging and analyses

All images were captured using an Olympus IX73 inverted fluorescence microscope linked to a PCO/Cooke Corporation Sensicam QE cooled CCD camera. An UPlanSApo 100x objective lens (oil) with a 1.40 numerical aperture was used. A filter wheel system with GFP/mCherry-ET Sputtered series with high transmission (Biovision Technologies) was used. For all images, cells were grown in the LabTek Chambered #1.0 borosilicate coverglass system (Nalge Nunc International, Rochester, NY). The IPLab software was used for image acquisition and analysis. Image labeling was done using Microsoft PowerPoint and/or Adobe Photoshop. For quantitation of dynein signal intensity, a region of interest (ROI) was selected and the Max/Min tool of the IPLab program was used to measure the maximal intensity within the ROI. The ROI box was then dragged to a nearby area outside of the cell to take the background value, which was then subtracted from the intensity value. For quantitation of HookA-GFP-marked early endosome accumulation at the hyphal tip, the total signals at the hyphal tip (Sum) was measured, and background subtracted the same way. Hyphae were chosen randomly from images acquired under the same experimental conditions. For measuring the signal intensity of a MT plus-end comet formed by GFP-dynein, only the comet close to hyphal tip was measured. For measuring GFP-dynein signal intensity at septa, usually only the septum most proximal to the hyphal tip was measured. Images were taken at room temperature immediately after the cells were taken out of the incubators. Cells were cultured overnight in minimal medium with 1% glycerol and supplements at 32°C.

### Statistical analysis

All statistical analyses were done using GraphPad Prism 9 for macOS (version 9.2.0, 2021). The D’Agostino & Pearson normality test was performed on all data sets. For all data sets, non-parametric tests were used without assuming Gaussian distribution, except that Ordinary one-way ANOVa was used for the analysis presented in Supplemental Figure 2B. Specifically, the Kruskal-Wallis ANOVA test (unpaired) with Dunn’s multiple comparisons test was used for analyzing multiple data sets, and the Mann-Whitney test (unpaired, two tailed) was used for analyzing two data sets. Note that adjusted p values were generated from the Kruskal-Wallis ANOVA test with Dunn’s multiple comparisons test. In all figures, **** indicates p<0.0001, *** indicates p<0.001, ** indicates p<0.01, * indicates p<0.05 *, and “ns” indicates p>0.05.

## Acknowledgements

We thank Berl Oakley, Elizabeth Oakley, Miguel Peñalva and Reinhard Fischer for Aspergillus strains. This work was funded by the National Institutes of Health RO1 GM121850 (to X. Xiang) and R35 GM140792 (to X. Xiang). Disclaimer: The opinions and assertions expressed herein are those of the author(s) and do not necessarily reflect the official policy or position of the Uniformed Services University or the Department of Defense.

## Author contributions

R.Q. and X.X. conceived the experiments. R.Q., J.Z. and X.X. carried out the experiments. X.X. and R.Q. wrote the manuscript, which was read and edited by all authors.

## Supplemental Materials

**Supplemental Table 1.**
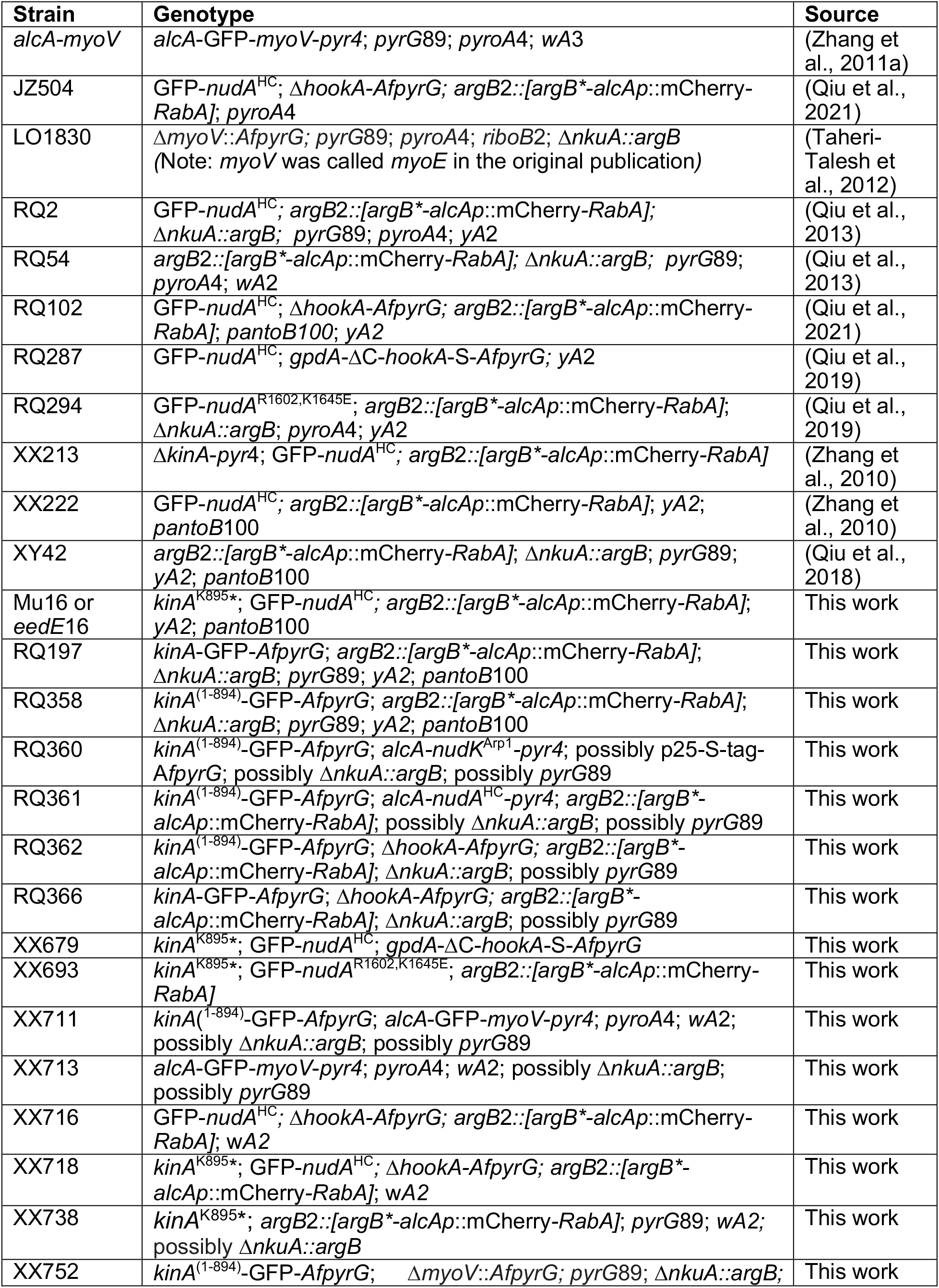
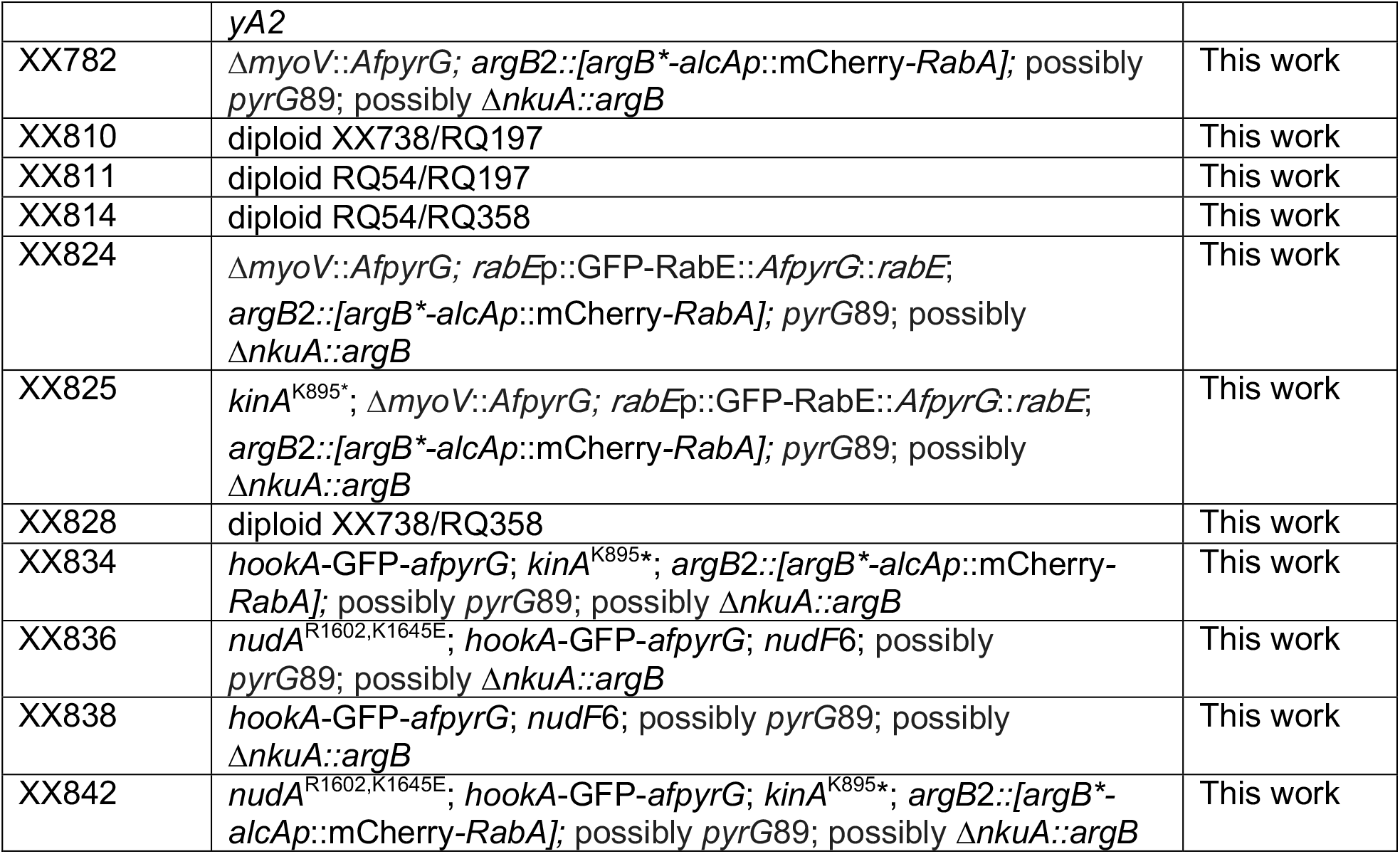
*Aspergillus nidulans* strains used in this study.

**Supplemental Figure 1.**
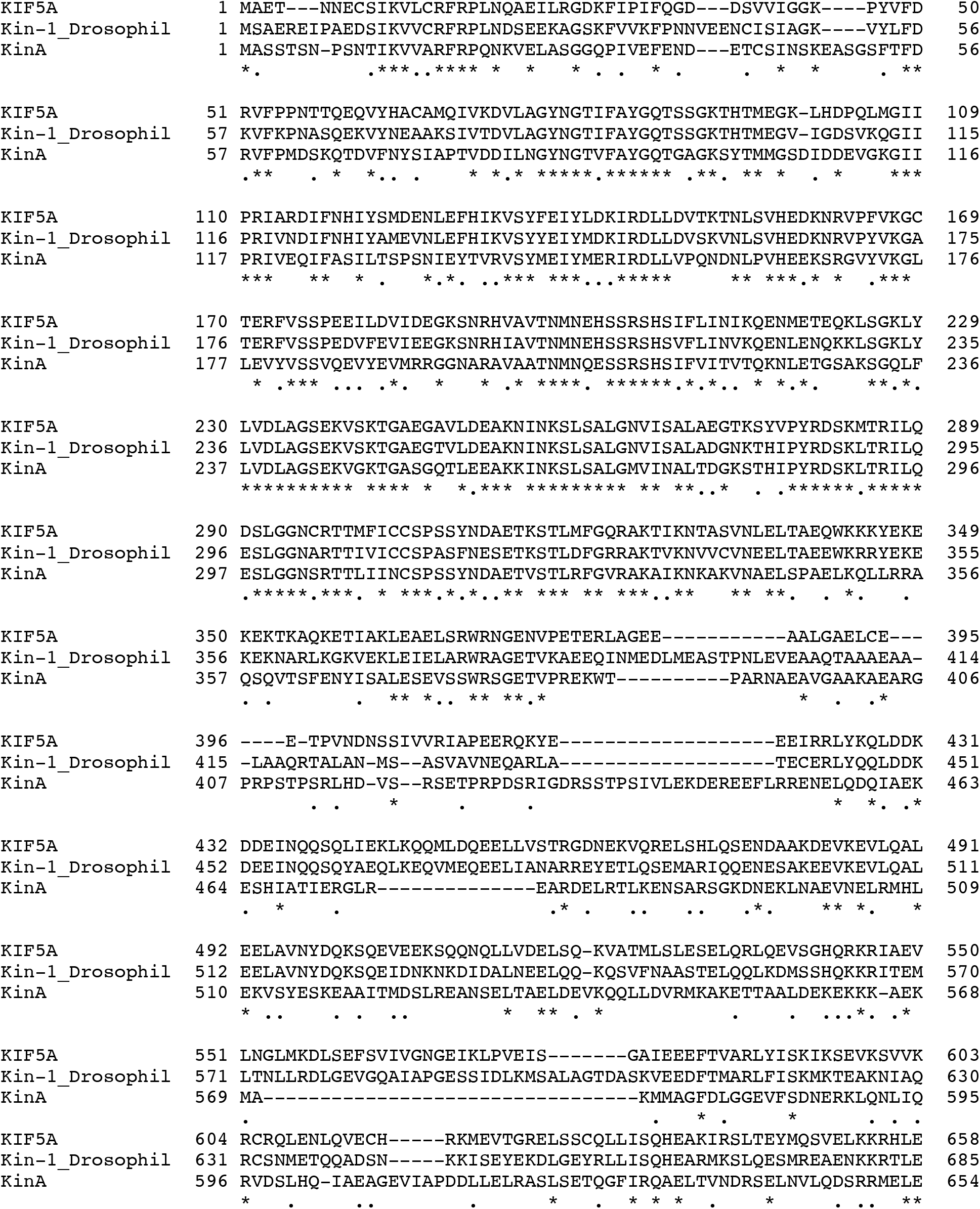
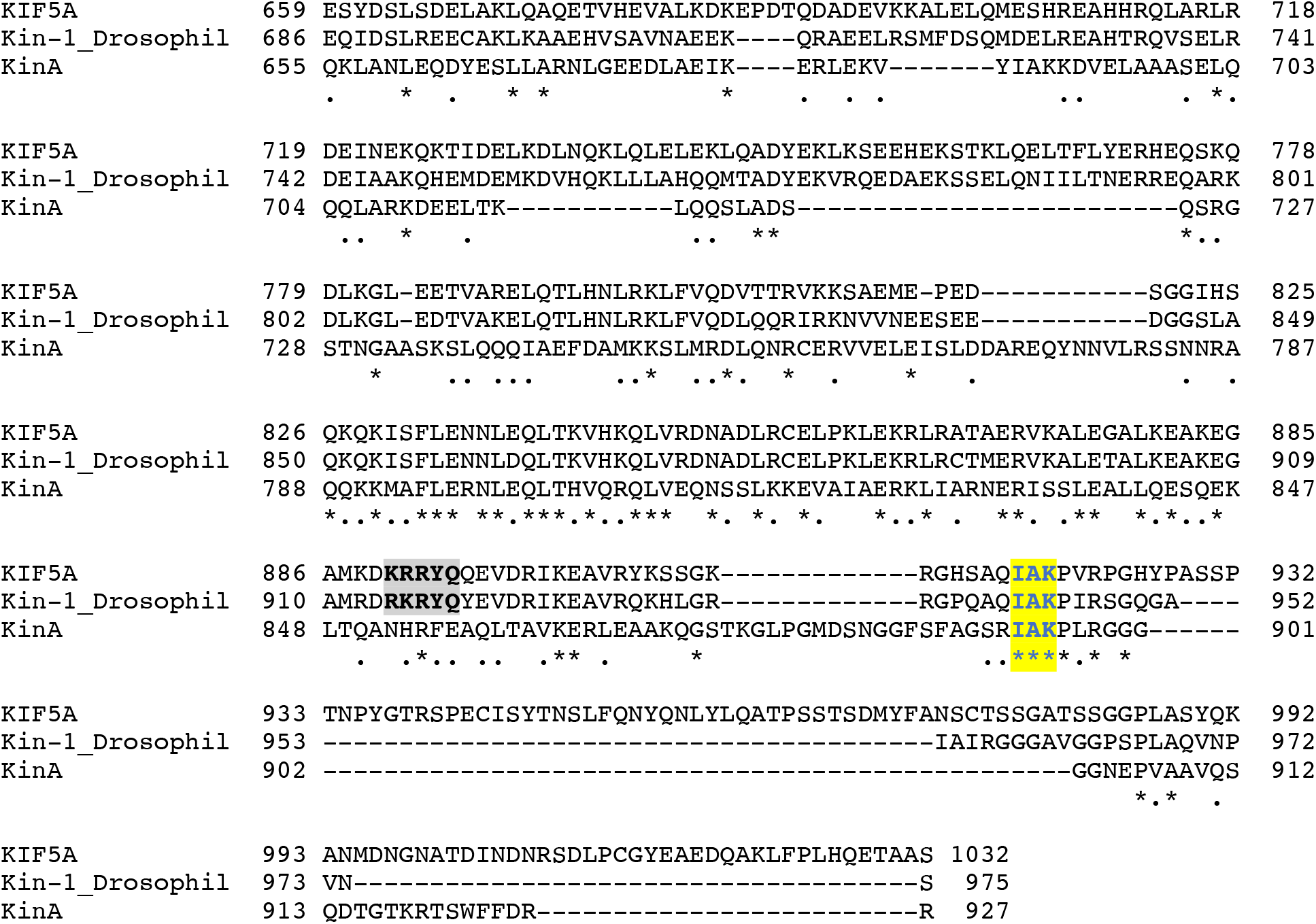
A sequence alignment of kinesin-1s including KIF5A, kinesin-1 heavy chain in Drosophila (Kin-1_Drosophil) and kinesin-1 in *Aspergillus nidulans* (KinA). The alignment was done using MacVector T-Coffee multiple sequence alignment (Pairwise Mode: Myers Miller). Residues that are identical (*), strongly similar (:) or weakly similar (.) are indicated. The IAK motif was highlighted in yellow and the C-terminal microtubule-binding domain was shaded with grey.

**Supplemental Figure 2.**
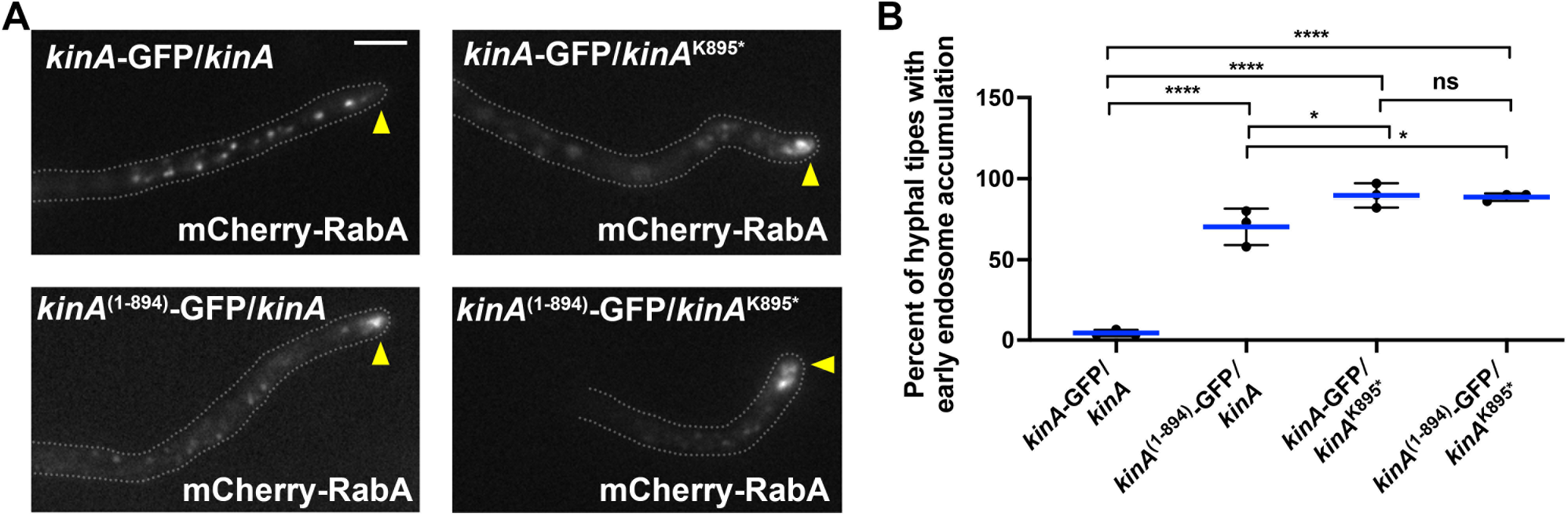
The *kinA*^k895*^ or the *kinA*^(1-894)^-GFP allele causes a defect in dynein-mediated early endosome transport in diploids with the presence of full-length *kinA*. (A) Microscopic images showing the distributions of mCherry-RabA-labeled early endosomes in four different diploids. Hyphal tip is indicated by a yellow arrowhead. Bar, 5 μm. (B) A quantitative analysis on the percentage of hyphal tips with the abnormal accumulation of early endosomes. Three experiments were performed, and in each experiment, 30 or more hyphal tips were examined for each strain. Scatter plots with mean and S.D. values were generated by Prism 9. ****p<0.0001; *p<0.05; ns, non-significant or p>0.05 (Ordinary one-way ANOVA, unpaired).

